# Divergent genotype-phenotype maps determine the capacity for adaptation to antibiotics among Enterobacteria

**DOI:** 10.64898/2026.10.06.756972

**Authors:** Rhea Vinchhi, Ashani Joshi, Seona Paranjape, Raghav Galgali, Chetna Yelpure, Shraddha Karve, Nishad Matange

**Author notes:** Equal contribution (Ashani Joshi, Seona Paranjape).

## Abstract

What dictates the extent to which a bacterial strain can evolve under antibiotic pressure remains poorly defined, despite its implications for the effectiveness of antibiotics in clinical practice. In this study, we compare the evolution of *Escherichia coli* and *Klebsiella aerogenes,* two medically relevant members of *Enterobacteriaceae*, under colistin and trimethoprim selection. We discover that the mere presence of effector genes of resistance, as well as conserved gene regulatory architecture, do not necessarily translate to similar evolvabilities during antibiotic challenge. Instead genotype-phenotype maps of adaptive mutations, which can diverge at different biological levels, govern how effectively a strain evolves resistance. Both, *E. coli* and *K. aerogenes*, demonstrated strikingly parallel capacities for adaptation to trimethoprim, as a result of conserved mechanisms of resistance, including drug target modification and efflux. On the other hand, *E. coli* was driven to extinction by colistin, while *K. aerogenes* was able to evolve a high level of colistin resistance. Like the better studied *K. pneumoniae*, *K. aerogenes* acquired mutations in the BasSR (also called PmrAB) and PhoQP two-component signalling pathways under colistin selection. These mutations increased the expression of chromosomally-encoded lipopolysaccharide (LPS) modifiers, EptA and Arn. Similar mutations in *E. coli* were unable to confer protection against colistin, despite successfully up-regulating homologous effectors genes. However, expression of MCR-1, a horizontally acquired EptA-homolog found in clinical strains, conferred high level colistin resistance to *E. coli*. This difference in the intrinsic capacity to adapt to colistin was reflected in frequencies of resistance among clinical isolates from around the world. It also altered the dependency of *Escherichia* and *Klebsiella* species on horizontal gene acquisition for colistin resistance. Our study highlights the importance of understanding pathogen-specific genotype-phenotype maps of drug resistance to better predict long-term effectiveness of antibiotic therapies and define resistance-proof treatment strategies.

## Introduction

Adaptation to environmental fluxes determines the evolutionary success of an organism and the capacity for adaptation, i.e. evolvability, defines its limit [1–4]. For bacteria, evolvability is measured in terms of the rate or extent of adaptation under specific environmental challenges [5–8], such as antibiotic exposure that results in antimicrobial resistance (AMR). Given severity of the AMR crisis [9], understanding what determines evolvability during antibiotic challenge is crucial to curb drug-resistant infections.

The evolution of antibiotic resistance has been studied using two approaches: laboratory evolution of *de novo* resistance [6, 8, 10–13] or studying drug-resistant strains directly in clinical or environmental contexts [14–17]. While the latter approach provides real world insight, the former is better suited for mechanistic investigation. For example, a recent metanalysis revealed that minimum inhibitory concentrations (MICs) of antibiotics for clinical isolates of different pathogens were correlated with complex factors, such as patient age and site of infection [18]. On the other hand, laboratory studies have been invaluable in teasing apart the role of individual environmental parameters such as strength of antibiotic selection [19–21], duration of exposure [22], nutrient availability [23, 24] and population bottleneck sizes [25]. A common finding from both approaches is the role of genetic constitution of bacteria as a determinant of evolvability under drug pressure [8, 10, 11, 13, 26–35]. Indeed, the pre-existence of genes encoding effectors of antibiotic resistance, such as genomically-encoded beta-lactamases [36] or multi-drug efflux pumps [11, 26], directly improves evolvability. Likewise, access to beneficial mutations in effector genes also determines the capacity for adaptation. For instance, evolvability may be limited when multiple small-effect mutations are required [37], whereas epistasis between beneficial mutations amplifies evolvability, especially at high concentrations of antibiotics [10, 38–40].

Effector genes of antibiotic resistance are frequently conserved between related bacterial species. This is exemplified by the conservation of homologous effector genes across prominent species belonging to the family Enterobacteriaceae (Supplementary Figure 1). The presence of similar effector genes in different members of this family would suggest similar capacity for resistance evolution as well. Here, we experimentally test this hypothesis using comparative laboratory evolution for two enterobacteria, *Escherichia coli* and *Klebsiella aerogenes*. *E. coli* is a gut commensal, though several strains are associated with multi-drug resistant infections of the intestine, urinary tract or bloodstream [41]. *K. aerogenes* is an emerging pathogen associated with increasing incidences of nosocomial infections of bloodstream, intestine, and urinary and respiratory tracts in humans [42, 43]. We challenged both species with two broad spectrum antibiotics relevant for Gram-negative infections, i.e. trimethoprim and colistin. Trimethoprim competitively inhibits bacterial Dihydrofolate Reductase (DHFR), encoded by the *folA* gene. Colistin binds to Lipid A in the lipopolysaccharide (LPS) layer of Gram-negative bacteria disrupting membrane integrity and causing cell death [44, 45]. Trimethoprim resistance is conferred by mutations at the drug-binding pocket of DHFR [30, 46–48], or by its transcriptional upregulation [13, 47, 49, 50]. Chromosomally-encoded effectors of colistin resistance include the Arn and EptA enzymes that transfer 4-amino-4-deoxy-L-arabinose (L-Ara4N) and phosphoethanolamine (PE), respectively, onto 1’-phosphate of Lipid A to reduce drug binding [44]. Colistin resistance requires overexpression of these effectors and is mediated by activating mutations in two-component signalling pathways PhoQP and BasSR (also called PmrBA) [44]. Apart from chromosomal mutations, enterobacteria also evolve colistin resistance via plasmid-borne *mcr* genes, which modify lipid A with PE at 4’-phosphate of lipid A [51].

We discover that *E. coli* and *K. aerogenes* demonstrate similar capacity for adaptation to trimethoprim, but differ dramatically when challenged with colistin. We explain these differences by systematically dissecting genotype-phenotype maps of adaptive mutations across the two species. Finally, we show that the intrinsic capacity for adaptation to antibiotics is also reflected in frequencies and mechanisms of resistance among clinical strains.

## Results

### Evolvability of two enterobacterial species under trimethoprim and colistin pressure

Laboratory strains of two enterobacterial species, *E. coli* K-12 MG1655 and *K. aerogenes* ATCC 13048, had similar susceptibilities to trimethoprim, colistin, rifampicin and streptomycin, among a panel of antibiotics (Figure 1A). Since MICs of trimethoprim and colistin (both ∼1 μg/mL) fell within the “susceptible” range based on CLSI breakpoints [52] and both species encoded similar chromosomal effectors of resistance (Supplementary Figure 1), we asked whether these bacteria also had similar capacity for resistance to the two antibiotics. To address this question, we employed laboratory evolution for five replicate populations each of *E. coli* and *K. aerogenes*. Serial passaging was performed for 30 cycles of growth (∼210 generations) under trimethoprim or colistin pressure such that 1% of the population was transferred to fresh growth media every 24 hours. In order to test the limit of adaptation, antibiotic concentration was increased 5-fold after every 5^th^ passage (∼35 generations), starting at 0.1x MIC and culminating at 62.5x MIC (Figure 1B).

**Figure 1.**
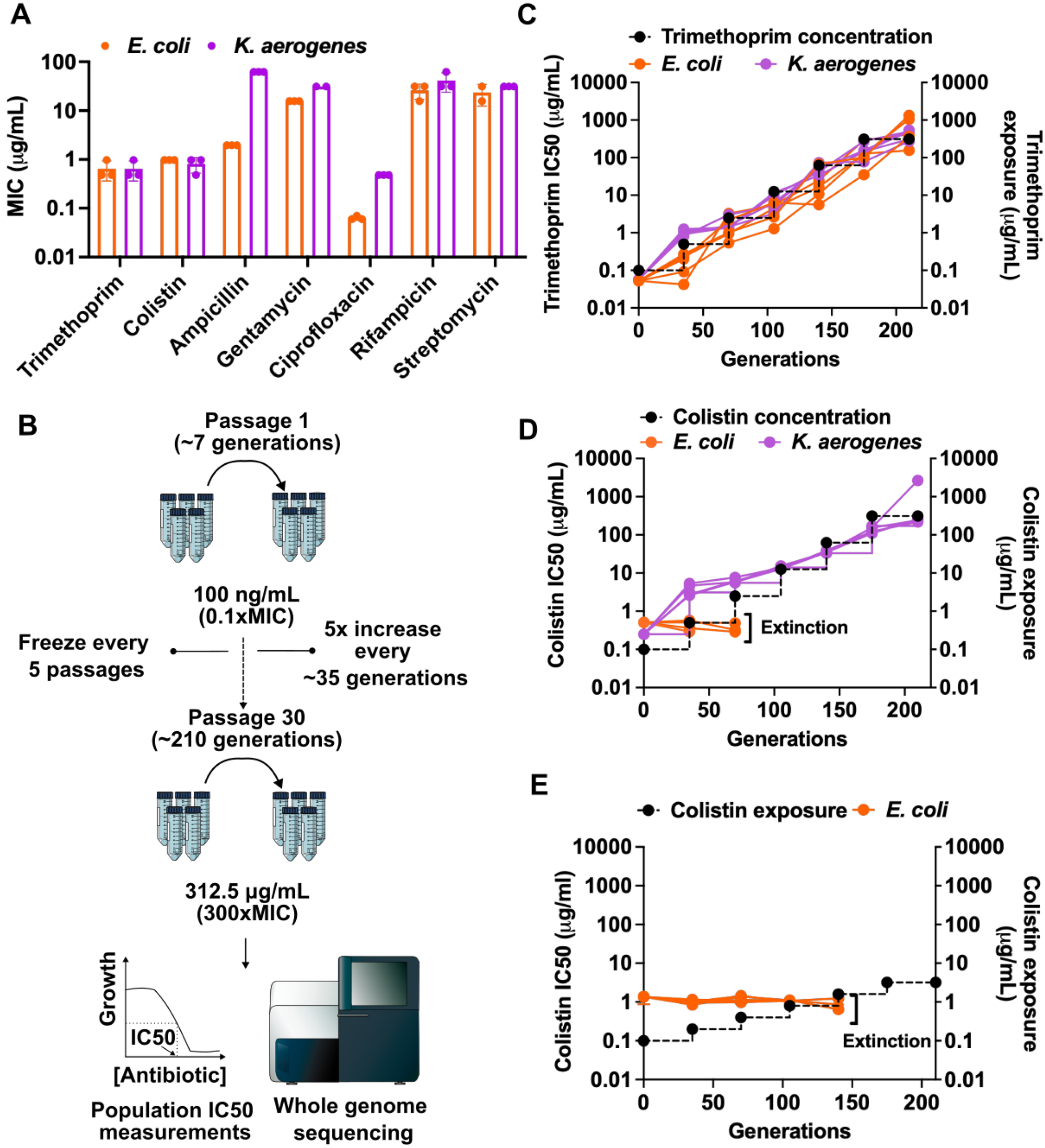
Comparing evolvabilities of *E. coli* K-12 MG1655 and *K. aerogenes* ATCC 13048 under trimethoprim and colistin pressure: **A**. MIC values of trimethoprim, colistin, ampicillin, gentamicin, ciprofloxacin, rifampicin and streptomycin for *E. coli* (orange) and *K. aerogenes* (purple) calculated from broth-dilution experiments. Mean ± SD of three replicates are plotted as bars and individual replicates are shown. **B**. Schematic of the comparative laboratory evolution used in this study. Five replicate lineages each of *E. coli* and *K. aerogenes* were evolved in LB media with increasing concentrations of trimethoprim or colistin for a total of 30 passages (∼210 generations). Each growth cycle, which lasted for 24 hours, corresponded to ∼7 generations. Antibiotic concentration was increased by 5-fold every ∼35 generations, starting at 100 ng/mL at Passage 1 and ending at 312.5 μg/mL at Passage 30. Aliquots from evolving populations were frozen after every 5 passages (∼35 generations) for further analysis, including population IC50 measurements and whole genome sequencing. **C, D.** Trimethoprim (**C**) and colistin (**D**) IC50s for *E. coli* (orange) and *K. aerogenes* (purple) populations evolving under antibiotic pressure are plotted on the left Y-axis. Mean of measurements performed in duplicate are shown. Each trace represents an individually evolving lineage. Black dotted line shows the antibiotic concentrations (in μg/mL on the right Y-axis) during evolution. The time points at which *E. coli* populations evolving under colistin pressure went extinct are indicated. **E**. Colistin IC50s of five independently evolving *E. coli* populations, in a less strict regime, where colistin concentration was doubled after every ∼35 generations. Black dotted line represents antibiotic exposure (in μg/mL on right Y-axis) during evolution.

Under trimethoprim pressure, all lineages of *E. coli* and *K. aerogenes* survived the full duration of the experiment, with IC50 values (concentration resulting in 50% growth inhibition) increasing in line with drug pressure (Figure 1C). When challenged with colistin, however, the two organisms showed divergent evolvability. *K. aerogenes* populations steadily increased colistin IC50 over evolution and were able to survive the highest antibiotic concentration tested. However, all 5 *E. coli* lineages were driven to extinction between 500 ng/mL (∼35^th^ generation) and 2500 ng/mL (∼70^th^ generation) of colistin, i.e. close to the MIC of the drug, without a detectable change in IC50 (Figure 1D). Even under less stringent selection regime, where colistin concentration was increased 2-fold every 35 generations, all *E. coli* populations went extinct between 800 ng/mL and 1600 ng/mL of antibiotic with undetectable changes in IC50 (Figure 1E). Thus, while the two bacteria had similar capacity for adaptation to trimethoprim, *E. coli* was unable to adapt as effectively as *K. aerogenes* under colistin pressure. This two antibiotic-two species system, therefore, provided an opportunity to investigate why evolvability under antibiotic challenge may diverge among enterobacteria despite the presence of conserved effector genes.

### Parallel molecular evolution of Dihydrofolate Reductase in *E. coli* and *K. aerogenes* during adaptation to trimethoprim

To understand the molecular basis for similar evolvability under trimethoprim pressure, we analysed genome sequences of *E. coli* and *K. aerogenes* at 35, 70, 105 and 210 generations of evolution. Point mutations in the *folA* gene and its promoter were observed in all lineages, demonstrating that drug-target modification was likely to be the primary mode of adaptation in both species. Two *E. coli* lineages additionally showed transient copy number changes at the *folA* locus that are known to augment resistance via DHFR overexpression [49] (Supplementary Figure 2). Consistent with earlier reports [48–50], increasing trimethoprim pressure resulted in sequential acquisition of *folA* mutations, such that multiple mutations were incorporated into DHFR at later time points in evolution (Figure 2 A,B, Supplementary File 1). Interestingly, strong parallelism was observed among mutations in DHFR between *E. coli* and *K. aerogenes* (Figure 2 A,B), reflecting the high conservation between orthologous enzymes (∼150-fold over wild type, Supplementary Figure 3). Further, specific combinations of mutations in DHFR also appeared repeatedly across lineages of the two organisms. Most prominent among them were D27E and F153S that co-occurred in 2 out of 5 lineages of both organisms (Figure 2A, B). Reconstructing D27E and F153S on a plasmid-borne copy of DHFR in *E. coli* confirmed positive epistasis between them (s = 46.32 ; σ_s_ = 41; Figure 2C, Supplementary File 2). Indeed, this double mutant conferred very high levels of trimethoprim resistance to *E. coli* (∼150-fold over wild type, Figure 2C), demonstrating that it was a “jackpot” combination and explained its repeated emergence during an increasing antibiotic selection regime. There was also a striking lack of co-occurrence of D27E or F153S with other mutations such as I94L or A26T/V, indicating additional epistatic constraints on mutational trajectories of DHFR (Figure 2A,B, Supplementary File 1). In support of this idea, mutation reconstruction experiments showed that most combinations of DHFR mutations that did not spontaneously co-occur tended to be less-than-additive (Figure 2C). In fact. some combinations showed strong negative sign epistasis, such as the combination of D27E or F153S with I94L (s < 0, Figure 2C, Supplementary File 2), explaining the early commitment to specific mutational trajectories. Earlier work has shown that negative epistasis between DHFR mutations occurs due to trade-offs with folding stability, and clearance from the bacterial cell by Lon protease limits the benefit of certain mutation combinations [30, 46]. Here too, we found that the benefit of D27E/F153S + I94L mutations was limited by Lon, as both combinations conferred a higher level of resistance in Lon-deficient *E. coli* than in wild type bacteria (Supplementary Figure 4). Thus, the strongly parallel molecular evolution of DHFR among *E. coli* and *K. aerogenes* was explained by epistasis-based constraints, and contributed to similar trajectories of genotypic and phenotypic evolution of trimethoprim resistance.

**Figure 2.**
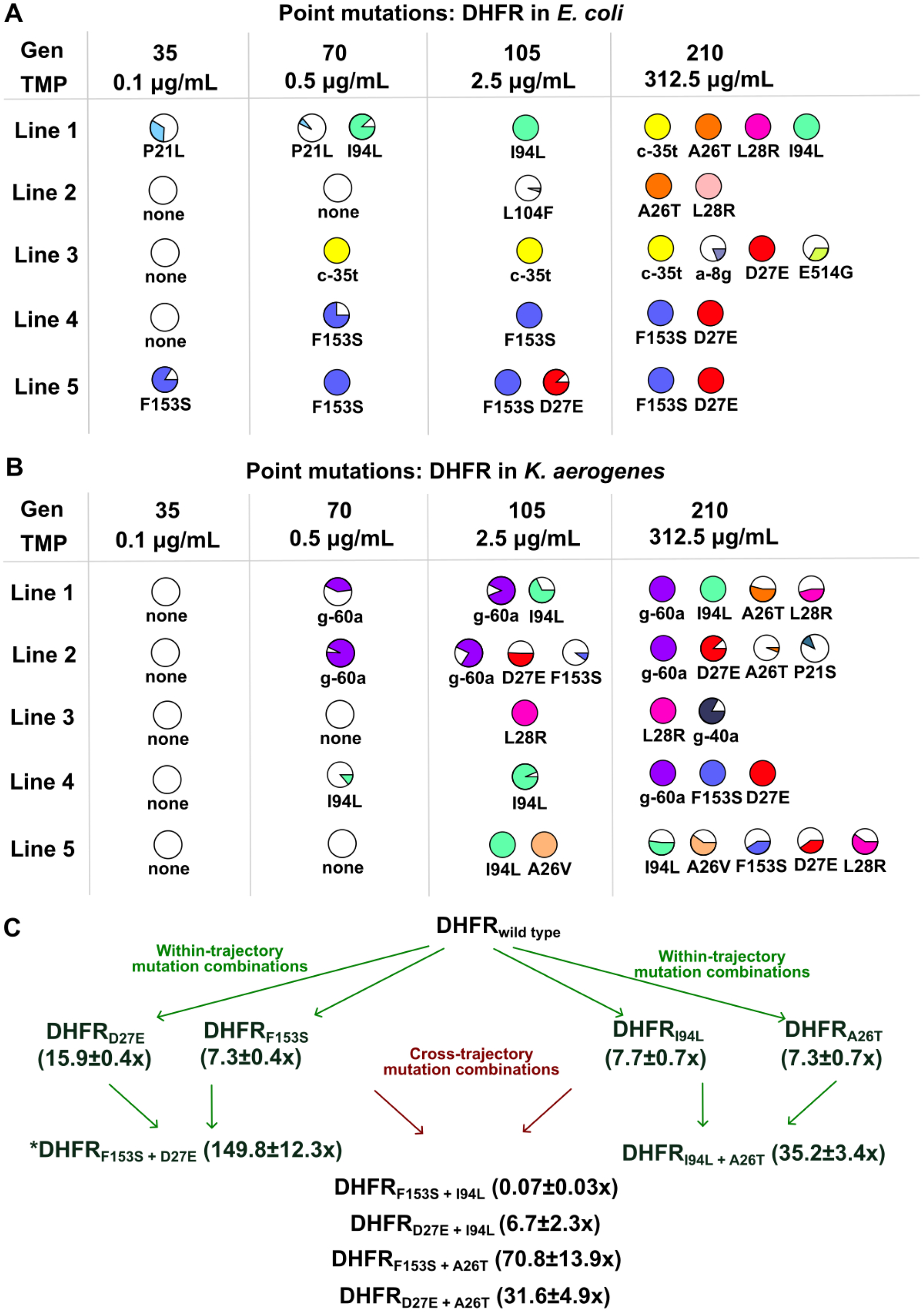
Similar mutation trajectories at the *folA* locus, coding for the target of trimethoprim, in *E. coli* and *K. aerogenes*. **A, B.** Frequency of promoter and coding mutations in *folA* (which codes for DHFR) in trimethoprim-adapted *E. coli* (**A**) and *K. aerogenes* (**B**) populations at different time points of evolution. Each mutation is represented as a pie for each lineage, where a complete circle indicates 100% frequency as determined by genome sequencing. For coding region mutations, amino acid substitutions are indicated by single letter amino acid codes in upper case. Mutations in the promoter are represented in lower case. Absence of mutations in DHFR is shown as an empty circle. **C**. Epistasis among DHFR coding-region mutations in *E. coli*, analysed by generating single and double mutations on a plasmid-borne copy of *folA.* Trimethoprim IC50s were measured for each mutant and expressed as fold over IC50 of a strain expressing wild type *folA* (set to 1). Mean ± SD of fold-IC50 values from three independent measurements for each mutant are shown in parentheses. Green arrows represent mutation combinations that were observed in the laboratory evolution experiment, i.e. within trajectory mutations combinations. Red arrows represent double mutation combinations which did not co-occur in any evolving lineage, i.e. cross-trajectory mutation combinations. The jackpot combination, DHFR_F153S + D27E_, which occurred repeatedly in both organisms is marked with an asterisk (*).

### Parallelism in evolution under trimethoprim pressure goes beyond mutations in DHFR alone

Earlier studies in *E. coli* [47, 48], and the above analyses suggested that sequential accumulation of mutations in DHFR could explain the high capacity for evolution of trimethoprim resistance in enterobacteria. We tested the validity of this model by reconstructing one of the observed mutational trajectories at the *folA* locus of *E. coli* (Lineage 5) and comparing IC50s of reconstructed and evolved genotypes. Lineage 5 was specifically chosen since it had fixed the jackpot D27E+F153S combination in DHFR (Figure 2). Scarless genomic knock-ins of *folA*-F153S and *folA*-D27E+F153S were created to mimic early and late time points, respectively. Surprisingly, though mutations in *folA* could explain 50-100 % of trimethoprim resistance that was observed at early time points, at the final time point, *folA* mutations alone explained only ∼1% of the observed IC50 values (Figure 3A). Thus, though target modification was the major contributor to resistance early during evolution, other adaptive strategies shaped the resistant phenotype further on in evolution.

**Figure 3.**
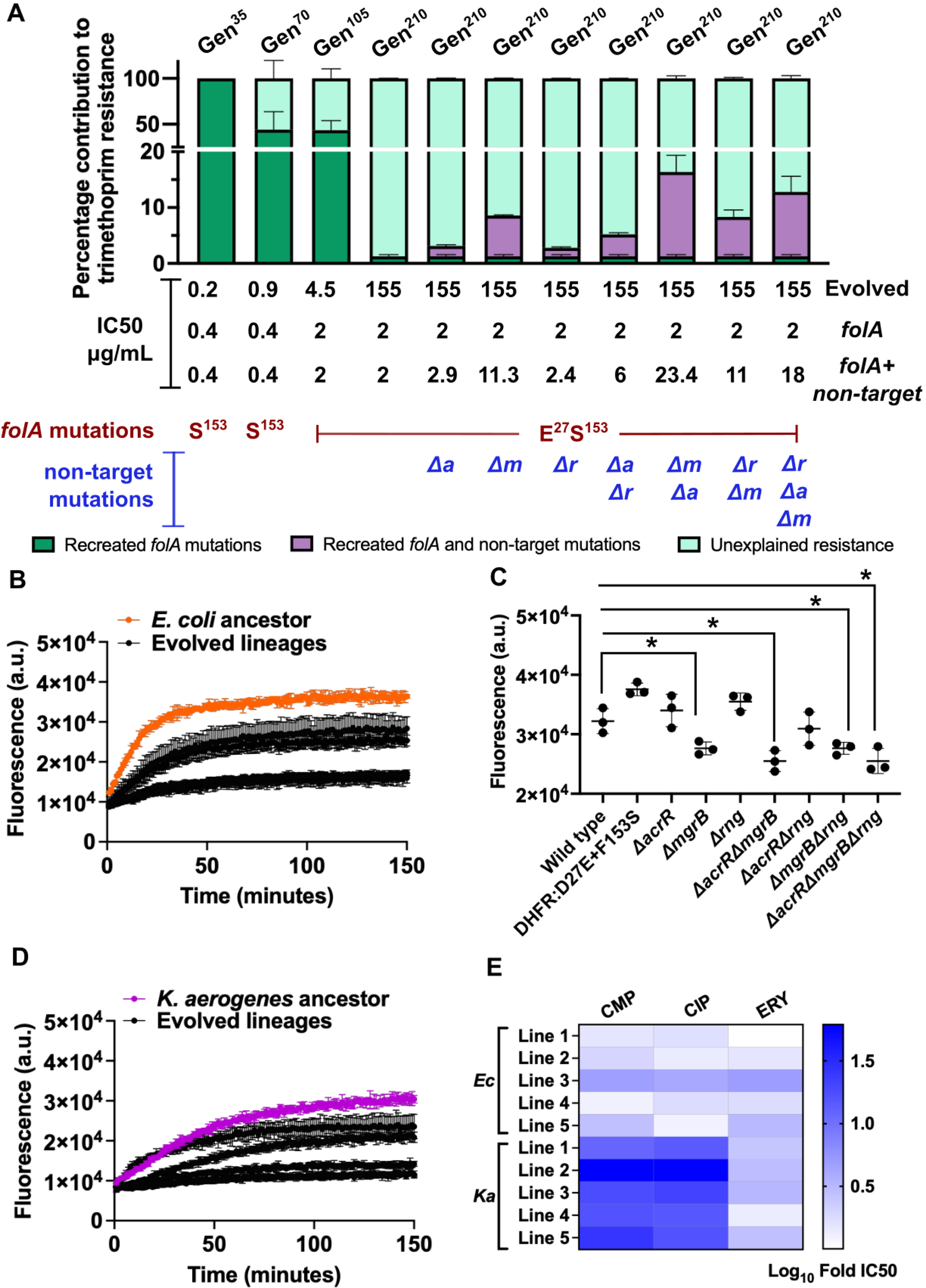
DHFR mutations alone are insufficient in explaining high trimethoprim resistance of evolving populations. **A.** Percentage contributions of *folA* and non-target mutations to trimethoprim IC50s of *E. coli* Line 5 are represented here. Trimethoprim IC50s of genomic knock-in *E. coli* strains harbouring *folA* F153S (S^153^) or *folA* F153S+D27E (E^27^S^153^)have been compared with the evolving Line 5 at time points indicated above the bars (for instance, Gen^35^ corresponds to generation 35). Mean ±SD of percentage contribution from three independent measurements for each strain is plotted as a dark green bar. Three “non-target” gene knockouts (*ΔmgrB:Δm, ΔacrR:Δa and Δrng:Δr*) were introduced singly, in pairs and all together in the *E. coli folA* D27E+F153S background. Mean ± SD of percentage contribution of *folA* and non-target mutations, together, is shown as a purple bar. Remaining trimethoprim resistance, after considering contribution of *folA* D27E+F153S and non-target mutations, is labelled as unexplained resistance, and shown as a cyan bar. The value of trimethoprim IC50 (mean of three independent measurements) for each recreated mutant along with genotype, as well as IC50 of the evolved population are given the appropriate bar. For individual IC50 values of knock-in strains, please see Supplementary file 2. **B, C, D.** Efflux in trimethoprim resistant *E. coli* (**B**) and *K. aerogenes* (**D)** evolved populations at the final time point (∼210 generations) was assessed by the accumulation of Hoechst 33342 dye. Fluorescence (excitation: 355 nm and emission: 460 nm) was measured in intervals 150 minutes (in 1 minute steps) for log phase cultures incubated with 10 μM of Hoechst 33342 dye. Mean ± SD of fluorescence (a.u. : arbitrary units) for three biological replicates of ancestor (in orange/purple) and evolved populations (in black) are plotted here. Efflux of single, double and triple gene knockouts of non-target genes in *E. coli* at 150 minutes are plotted in (**C**). Statistical significance was tested using t-test with Welch correction. Strains with significantly lower Hoechst 33342 fluorescence (i.e. higher efflux) compared to wild type are shown by * (p-value < 0.05) **E**. Cross resistance of final time-point trimethoprim-adapted populations of *E. coli* and *K. aerogenes* to chloramphenicol, ciprofloxacin and erythromycin. IC50 values for each evolved population was normalised to appropriate ancestor, such that a fold change of 1 implied no change. Mean cross resistance was calculated from three replicates, log_10_ transformed, and represented as a single colour gradient. Individual IC50 values of evolved populations and ancestral strains are tabulated in Supplementary file 2

We asked if adding mutations in “non-target” genes, i.e. other than *folA*, that occurred later in evolution could better quantitatively recapitulate resistance. For *E. coli*, these mutations included loss of function in *mgrB*, which confers low-level resistance by elevating DHFR expression [13, 50], and *acrR*, which codes for the repressor of *acrAB* efflux pump that determines intrinsic trimethoprim susceptibility [26]. Mutations in *acrR* were observed in some of the trimethoprim-adapted *K. aerogenes* lineages as well (Supplementary File 1). A third locus, *rng* coding for RNase G (Supplementary File 1, Supplementary Figure 5), which when deleted confers resistance to trimethoprim (Supplementary Figure 3) and came up in an earlier screen for regulators of intrinsic resistance [26], was also included. All three genes had accumulated mutations that were partial or complete loss-of-function (Supplementary File 1, Supplementary Figure 3), allowing us to use gene knockouts as surrogates for the spontaneous mutants. Using knockouts of *mgrB*, *acrR* and *rng*, either singly or in combination, we assessed how IC50 values of knock-in strains harbouring DHFR mutations were altered. Among single gene deletions, only Δ*mgrB* increased trimethoprim IC50 of the *folA*-D27E+F153S knock-in substantially, i.e. by ∼5-fold (Figure 3A, Supplementary File 2). However, all three double knockouts and the triple knockout of non-target genes enhanced trimethoprim IC50 and could explain upto ∼11% of the resistance level observed at 210 generations (Figure 3A, Supplementary File 2). Notably, a majority of resistance was still left unexplained either due to other non-target mutations that we did not incorporate or phenotypic adaptations, such as transcriptional [53] or metabolic changes [54] induced by trimethoprim that would not be represented in the genome sequence.

To investigate the possible molecular mechanisms underlying target-independent adaptation we analysed transcriptomes of two of the non-target mutants. RNA sequencing analysis of the *mgrB* [13, 49] and *rng* (Supplementary File 3) knockouts showed that they independently resulted in mild up-regulation of efflux proteins (Supplementary Figure 5). Using intracellular accumulation of Hoechst H33342 dye as a readout, we observed increased efflux in *E. coli ΔmgrB*, *ΔmgrBΔacrR*, *ΔmgrBΔrng*, and *ΔacrRΔmgrBΔrng*, as well as all replicate populations of trimethoprim-evolved *E. coli,* corroborating our transcriptomics results (Figure 3B,C). Importantly, a high efflux phenotype was observed even among those *E. coli* lineages that did not harbour any of the 3 non-target mutations (Figure 3B), suggesting that other underlying mechanisms, including phenotypic adaptation, may be at play as well. Interestingly, all lineages of trimethoprim-adapted *K. aerogenes* also demonstrated increased efflux of Hoechst H33342 dye (Figure 3D). Further, we also observed cross resistance to chloramphenicol, ciprofloxacin or erythromycin, known substrates for efflux pumps [55–57], in trimethoprim-evolved *K. aerogenes* and *E. coli* populations (Figure 3E, Supplementary File 2), substantiating their high efflux phenotypes. Thus, increased efflux activity potentiated the resistance conferred by drug-target mutations and was required to quantitatively explain the resistance of evolved populations. Importantly, like drug-target mutations, these phenotypes were also conserved between *E. coli* and *K. aerogenes*. Taken together, our experiments showed that similar genotypic and phenotypic adaptive trajectories explained parallelism and similar evolvability of *E. coli* and *K. aerogenes* under trimethoprim pressure.

### *K. aerogenes* adapts to colistin by overexpressing chromosomal effectors of resistance

We next investigated why the capacity to evolve colistin resistance differed dramatically between *E. coli* and *K. aerogenes* (Figure 1). Though colistin had similar MIC for both species, there were differences in its bactericidal effects on ancestral *E. coli* and *K. aerogenes* cultures. While concentrations higher than MIC sterilised *E. coli* cultures, a small fraction of *K. aerogenes* (∼0.01%) cells remained viable even at 8x MIC (Figure 4A), indicating that the latter was more likely to survive high drug pressure. Additionally, though colistin treatment induced similar level of outer membrane damage in both species (measured by NPN permeabilization [45]), inner membrane damage (measured by PI permeabilization [45]) was far greater for *E. coli* (Figure 4B, 4C). These experiments provided a mechanistically explanation for different intrinsic susceptibility at >MIC colistin. We also observed well-skipping and within-population heterogeneity during MIC measurements for *K. aerogenes* (Supplementary Figure 6) possibly hinting at a pre-existing phenotypically resistant sub-population that was absent in *E. coli*.

**Figure 4.**
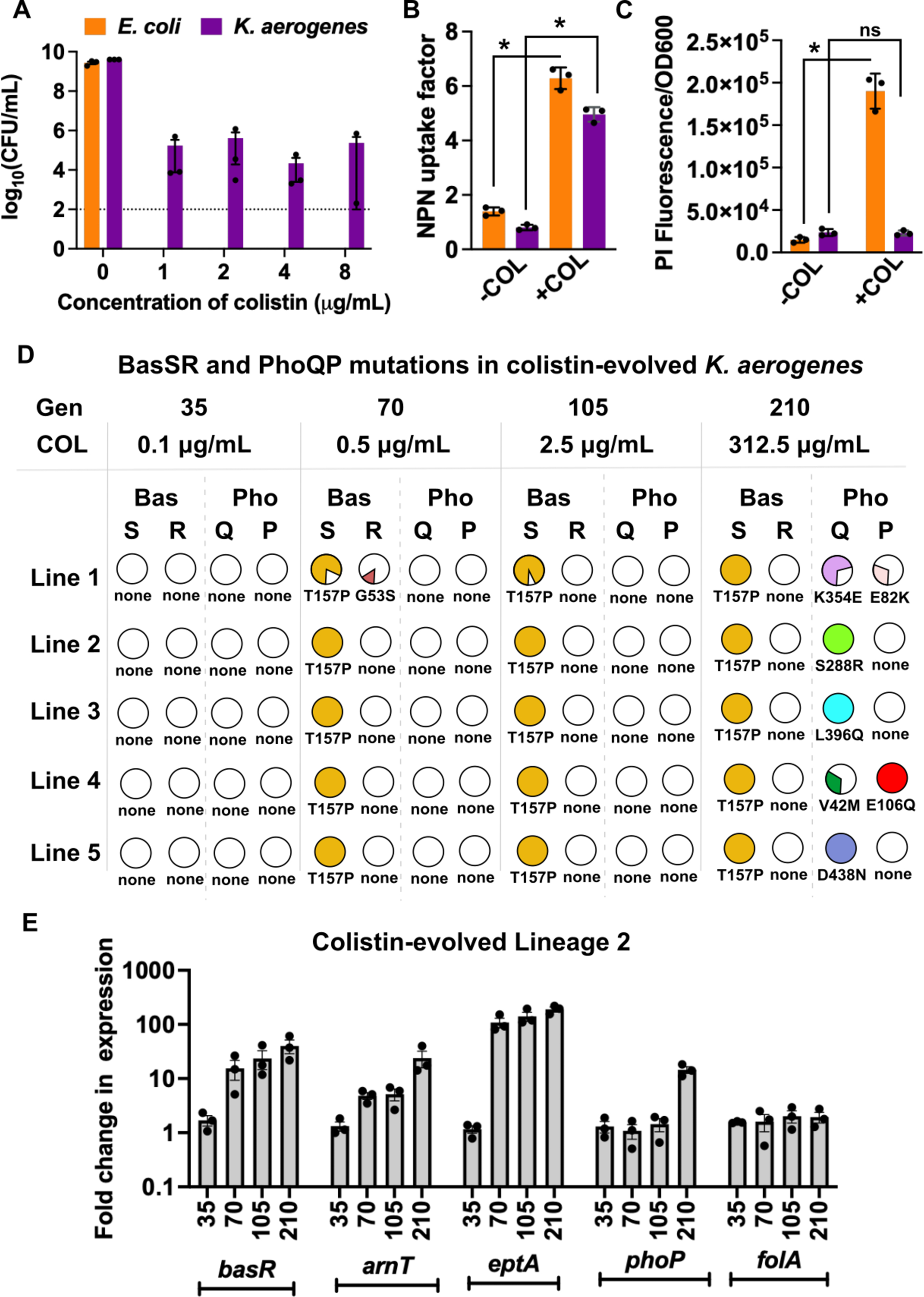
Genetic basis of colistin adaptation in *K. aerogenes*. **A**. Bactericidal activity of colistin, measured as viable counts (log_10_CFU/mL) of *E. coli* (orange) and *K. aerogenes* (purple) ancestral strains after 6 hours of growth in 0, 1, 2, 4 and 8 μg/mL colistin. Detection limit of the assay (∼100 CFU/mL) is indicated by a dotted line. No detectable growth was observed for *E. coli* above 1 μg/mL of colistin. Mean ±SD from three replicates is plotted as bars and individual data points are overlaid. **B.** Outer membrane permeabilization, quantified by NPN uptake factor (increases with increased membrane damage, see methods for details), for ancestral *E. coli* (orange) and *K. aerogenes* (purple) without colistin treatment (-COL) and upon exposure to 1 μg/mL colistin (+COL). Mean ±SD of three replicates is plotted as bars and individual data points are overlaid. **C.** Inner membrane permeabilization, quantified using propidium iodide fluorescence (which increases with membrane damage) for ancestral *E. coli* (orange) and *K. aerogenes* (purple) in the absence of colistin (-COL) or upon exposure to 1 μg/mL colistin (+COL). Mean ± SD of ratios of PI fluorescence and optical density at 600 nm from three replicates are plotted as bars and individual data points are overlaid. Statistical significance was tested using t-test with Welch correction and p<0.05 is indicated by *. **D.** Mutations in *basS*, *basR*, *phoQ* and *phoP* genes in *K. aerogenes* lineages at different generations (Gen) of evolution under colistin (COL) pressure. Mutations are indicated as single letter amino acid codes. Frequencies of mutations in the evolving population is represented as a pie, such that a filled circle represents 100%. **B**. Expression levels of *basR*, *arnT*, *eptA*, *phoP* and *folA* transcripts were measured by quantitative real-time PCRs in *K. aerogenes* Line 2 at 35, 70, 105 and 210 generations of evolution (indicated below individual bars). Data shown are expressed as fold change in expression compared to the *K. aerogenes* ancestor (set to 1). Mean ± SEM from three biological replicates are plotted as bars, and individual data points are shown.

Since *K. aerogenes* is poorly studied, we first identified adaptive mutations in this species under colistin pressure using genome sequencing, and compared them with what has been observed for the better-studied *K. pneumoniae* [44]. At 35 generations of evolution, no consistent mutational pattern was discernible (Supplementary File 1), despite a ∼15-fold increase in MIC (Figure 1D), indicating that early adaptation may be mediated by several low-frequency small-effect mutations. Subsequently, all 5 lineages first acquired the *basS*-T157P mutation, followed by mutations in *phoQ* (K354E, S288R, L396Q, V42M, D438N) (Figure 4D, Supplementary File 1). BasS-T157P and some of the observed *phoQ* mutations have been reported to confer colistin resistance in clinical strains of *K. pneumoniae* [58–65] by increasing expression of *arn* operon and *eptA* [44] (Figure 5A). We verified that *arnT* and *eptA* were indeed up-regulated in one of the evolving replicates of *K. aerogenes* (Lineage 2) using qRT-PCR (Figure 4E). These analyses confirmed that genetic mechanisms of high level colistin resistance were similar between *K. aerogenes* and *K. pneumoniae* and involved transcriptional up-regulation of Arn and EptA enzymes by mutations in signalling proteins.

**Figure 5.**
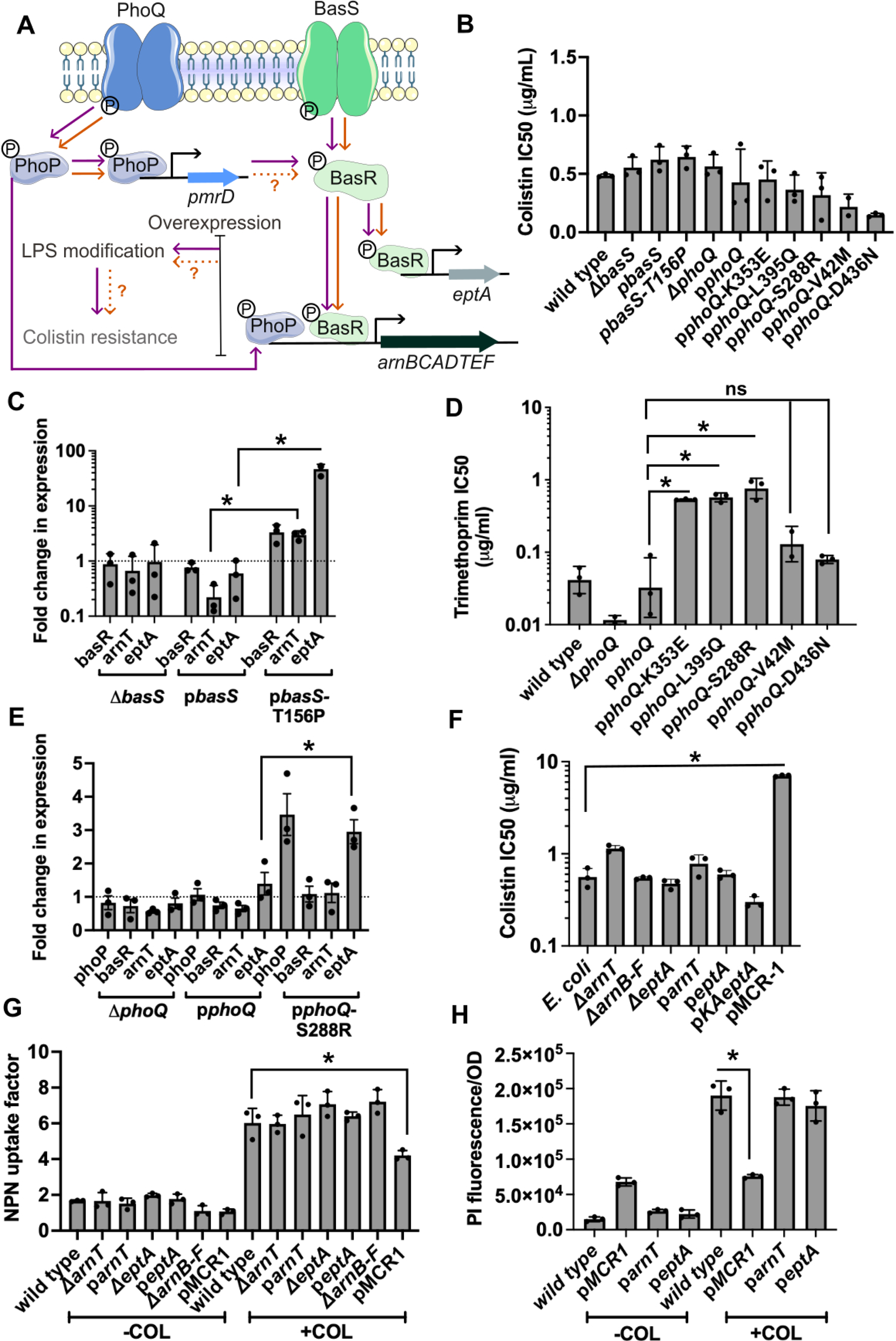
Overexpression of chromosomal effectors of resistance in *E. coli* is unable to protect it from colistin-mediated membrane damage. **A.** Schematic representation summarizing existing literature [66–76] about the role of PhoQP and BasSR two-component systems in conferring colistin resistance in *E. coli* and *K. pneumoniae*. Orange arrows are for *E. coli* and purple arrows are for *K. pneumoniae*. Ambiguous steps of the pathway, for which conflicting information exists in literature, are assigned dotted arrows with question marks. **B**. Colistin IC50 values of *E. coli* strains heterologously expressing wild type and mutant *basS* and *phoQ*. These mutations were introduced in *E. coli* using the high copy plasmids pCA24N-*basS* (p*basS*) and pCA24N-*phoQ* (p*phoQ*). Mean ±SD of three replicates is plotted as bars and individual data points are shown. None of the strains showed significant increase in colistin IC50 compared to wild type (unpaired t-test, p>0.05). C. Expression levels of *basR*, *arnT*, *eptA* transcripts measured by quantitative real-time PCR in *E. coli* Δ*basS*, p*basS* and p*basS*-T156P are plotted as fold over *E. coli* wild type. Data represent mean of three replicates ± SEM. A value of 1 indicates no change in expression and is represented by a dotted line. **D**. Trimethoprim IC50 values of *E. coli* strains heterologously expressing mutant *phoQ* proteins. These mutations were identified in *K. aerogenes* colistin resistant lines and introduced in *E. coli* using a high copy plasmid (pCA24N-*phoQ*). Trimethoprim resistance was used as a proxy for PhoQP pathway activation. Mean ±SD of three replicates is plotted. Mutants K353E, L395Q and S288R showed significantly higher trimethoprim IC50 values than ancestor (*: unpaired t test, p<0.05). **E**. Expression levels of *phoP, basR*, *arnT*, *eptA* transcripts measured by quantitative real-time qPCR in *E. coli* Δ*phoQ*, p*phoQ* and p*phoQ*-S288R are expressed as fold change over wild type. Data represent mean of three replicates ± SEM. Value of 1 indicates no change and is indicated by a dotted line. **F**. Colistin IC50 values of *E. coli* strains harbouring indicated gene deletions or overexpressing effectors of colistin resistance. Mean ±SD of three replicates is plotted. *E. coli* expressing Mcr1 was the only strain that demonstrated significantly greater colistin resistance than wild type (*: unpaired t test, p<0.05). **G**. Outer membrane permeabilization quantified by NPN uptake factor, for *E. coli* strains harbouring indicated gene deletions and overexpressing effectors of colistin resistance, at 0 and 1 μg/mL colistin. Mean ±SD of three replicates is plotted. *E. coli* expressing Mcr1 was the only strain that demonstrated significantly greater colistin resistance than wild type (*: unpaired t test, p<0.05). **H**. Inner membrane permeabilization, quantified using propidium iodide fluorescence for *E. coli* strains harbouring indicated gene deletions and overexpressing effectors of colistin resistance. Mean of three replicates (±SD) is plotted. *E. coli* expressing Mcr1 was the only strain that demonstrated significantly greater colistin resistance than wild type (*: unpaired t test, p<0.05).

### Up-regulation of chromosomal effectors is insufficient to confer high level colistin resistance in *E. coli*

We next asked why *E. coli* was unable to adapt to colistin despite encoding homologs of all the genes implicated in *Klebsiella* (Supplementary Figure 1). There exist conflicting reports regarding the architecture of relevant regulatory pathways in *E. coli* [66–76] (Figure 5A). Therefore, we took an unbiased and step-wise approach to investigate this question. First, we confirmed that there were no mutations in BasSR/PhoQP systems in *E. coli* at the last surviving time point, i.e. Generation 35, using genome sequencing (Supplementary File 1). Second, we asked if *basS* and *phoQ* mutations observed in *K. aerogenes* could confer colistin resistance in *E. coli*. However, introduction of these mutations in *basS* and *phoQ* of *E. coli* using a heterologous expression system did not increase colistin IC50 (Figure 5B). Third, we asked if the lack of colistin resistance was because the PhoQ and BasS proteins of *E. coli* were recalcitrant to activation by these mutations. BasSR and PhoQP pathways are positively autoregulated, and therefore levels of *basR* and *phoP* transcripts are indicative of the activation status of these pathways. Using qRT-PCR, we observed that *basR* transcript was elevated ∼3.5-fold in *E. coli* upon expression of BasS*-*T156P (homologous to BasS-T157P from *K. aerogenes*), confirming activation of the BasSR signalling pathway (Figure 5C). Three of the five mutations in *phoQ*, i.e. K353E, L395Q and S288R, increased trimethoprim resistance of *E. coli* (Figure 5D), a surrogate for PhoQP activity [50]. Further, PhoQ-S288R led to higher expression of *phoP* (Figure 5E), confirming activation of the PhoQP signalling cascade by these mutations. Thus, similar mutations could activate the BasS and PhoQ kinases in both organisms, but their activation did not confer colistin resistance. We reasoned that this discrepancy could be because activation of BasSR and PhoQP may not increase *arnT* and *eptA* gene expression in *E. coli*. However, this hypothesis was also incorrect, as the levels of *eptA* transcript were elevated in *E. coli* expressing BasS-T156P and PhoQ-S288R (∼46-fold and ∼3-fold, respectively) and *arnT* levels were elevated in *E. coli* expressing BasS-T156P (∼3-fold; Figure 5C, 5E). Finally, heterologous expression of the effectors directly from a multi-copy plasmid was also unable to increase colistin IC50s (Figure 5F) or alter colistin-induced NPN and PI uptake (Figure 5G, 5H), showing that increased expression of chromosomal effectors could not confer colistin resistance in *E. coli*.

This unexpected result could have two mechanistic explanations. First, EptA and Arn enzymes from *E. coli* had intrinsically lower activity than their homologs from *K. aerogenes*. However, plasmid-expressed *K. aerogenes*-*eptA* also did not confer colistin resistance in *E. coli* (Figure 5F). Alternatively, modifications of LPS could not confer resistance altogether in *E. coli*, or the specific modification of LPS by chromosomal effectors was inadequate in protecting *E. coli* from colistin. To distinguish between these possibilities, we expressed another LPS modifying enzyme, MCR-1, which chemically alters a different position compared to EptA [51] in *E. coli*. Expression of MCR-1 resulted in robust enhancement of colistin IC50 in *E. coli* (∼12-fold), and prevented colistin-induced NPN and PI uptake (Figure 5F,G,H). Likewise, neither EptA nor ArnT overexpression altered IC50 of a related antibiotic, polymyxin B, in *E. coli,* but MCR-1 did (Supplementary Figure 7). Thus, the reduced intrinsic capacity for adaptation to colistin stemmed from feeble protection provided by chromosomally encoded effectors of resistance in *E. coli*, rather than an inability to overproduce them by mutations in signalling pathways.

### Other strains of *E. coli* also have compromised evolvability under colistin pressure

*E. coli* K-12 MG1655 is a laboratory strain that has lost its O-antigen and is expected to have an altered cell envelope structure [77, 78]. To check if its evolvability was restricted by these differences in cell envelope, and therefore specific to the laboratory strain, we performed similar colistin evolution experiments with two other *E. coli* strains, Cft073, a domesticated uropathogen, and an environmental isolate from a sewage-contaminated river in Pune (India), belonging to ST type 998 with O2:H6 serotype (named Isolate S34). Like MG1655, both strains of *E. coli* showed a lower capacity for adaptation than *K. aerogenes*, being driven to extinction between the 70^th^ and 105^th^ generation. However, unlike MG1655 both strains did showed an increase in colistin IC50 (∼10-fold over respective ancestor) before going extinct, indicating greater evolvability than the laboratory strain (Figure 6A, B). Genome sequencing of surviving lineages showed mutations in *basSR* and *mgrB* in both strains indicating a similar mechanism of adaptation as *K. aerogenes*/*K. pneumoniae* (Figure 6A,B, Supplementary File 1) but with limited fitness benefit. Thus, though other strains of *E. coli* demonstrated higher capacity for colistin resistance than MG1655, they were still impaired compared to *K. aerogenes*. These results pointed to a genus-level compromise in evolvability under colistin pressure.

**Figure 6.**
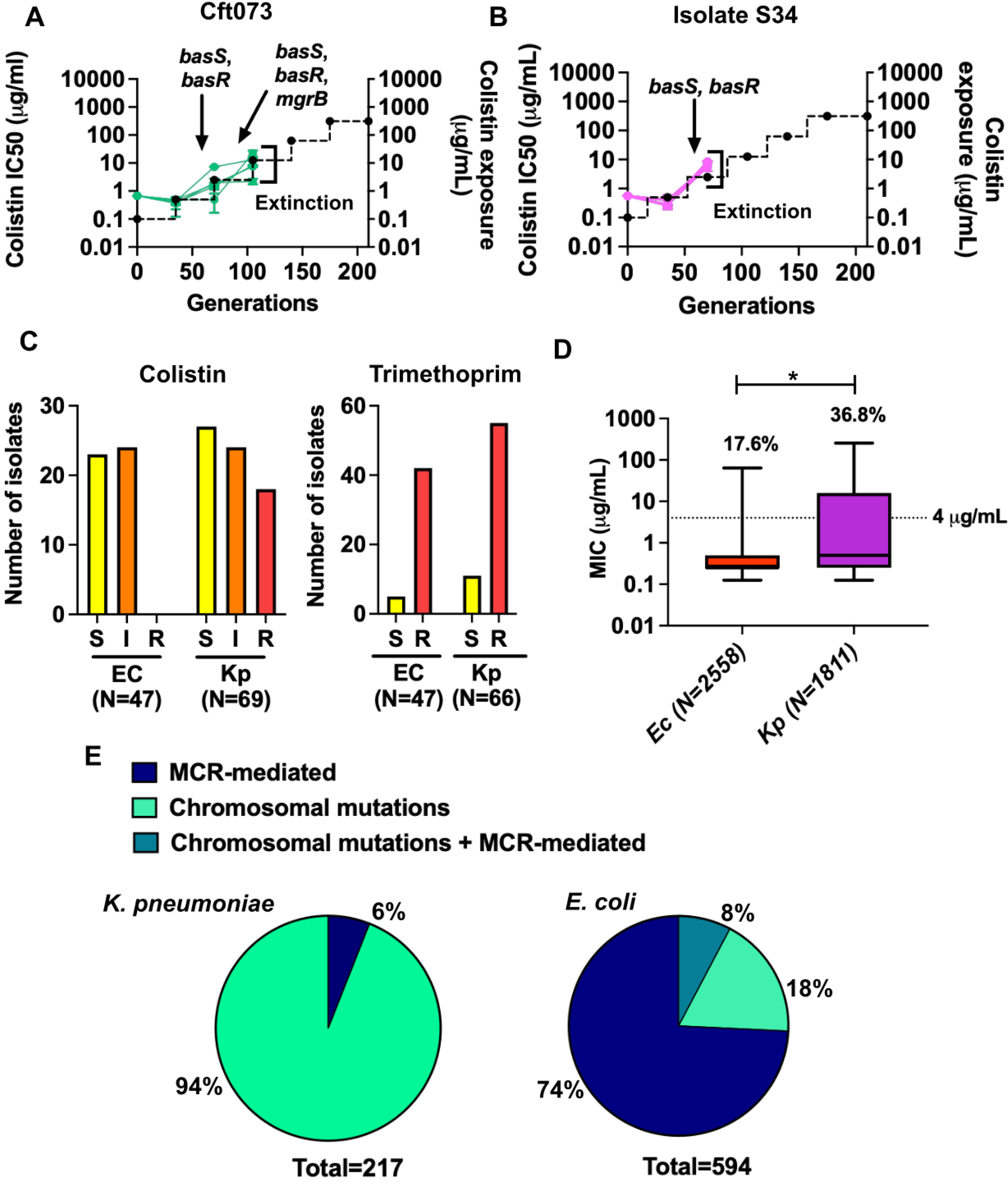
Lower intrinsic evolvability of colistin resistance in *E. coli* necessitates diverse mechanisms of resistance in clinical strains. **A, B.** Colistin IC50 values for five lineages of uropathogenic *E. coli* Cft073 (**A**) and environmental isolate *E. coli* S34 (**B**) evolving under colistin pressure are plotted. Each point indicates mean of two replicate measurements and each trace represents one of 5 replicate lineages. Black dotted line shows antibiotic exposure (in μg/mL, right Y-axis) during evolution, increasing by 5-fold every 35 generations. Extinction of the evolving populations is indicated in the graph. Mutations in the PhoQP and BasSR signalling pathways were detected in Cft073 (at 70 and 105 generations) and in S34 at (105 generations) as indicated. **C**. Colistin (panel on the left) and trimethoprim (panel on the right) sensitivity of hospital isolates of *E. coli* (N=47) and *K. pneumoniae* (N=66) from a recently published [79] data set from North and West India are plotted. Clinical isolates have been classified as sensitive (S, yellow bar), intermediate (I, orange bar) or resistant (R, red bar) according to CLSI guidelines. **D**. Distribution of colistin MICs (μg/mL) of clinical and veterinary isolates of *E. coli* (N=2558, orange bar) and *K. pneumoniae* (N=1811, purple bar) collated from 37 studies across 20 countries (details in Supplementary file 4) are plotted as box and whisker plots. Solid horizontal line within each box represents the median, and separates the second and third quartile of data. Whiskers represent the range of data. CLSI breakpoint for colistin resistance (4 μg/mL) is indicated with a dotted line. Percentage of strains of each species which have MICs above 4 μg/mL are indicated above each bar. Statistical significance was tested using an unpaired t-test with Welch correction (*: p-value < 0.05) E. Mechanisms of resistance associated with colistin resistant *K. pneumoniae* (N=217, left panel) and *E. coli* (N=594, right panel) isolates from around the world. Chromosomal mutations include PhoQP and BasSR loci. Percentage of isolates belonging to each category are noted in the pie charts as indicated.

### Intrinsic evolvability to colistin drives different mechanisms of resistance among clinical isolates of *E. coli* and *K. pneumoniae*

We investigated the real world implications of different evolvabilities, by analysing the frequencies of colistin resistance among a set of randomly sampled clinical isolates of *E. coli* and *K. pneumoniae* from New Delhi and Pune in India (Gheewala et al. 2026 [79]). Strikingly, close to 30% of *K. pneumoniae*, but none of the *E. coli* isolates, were resistant to colistin (i.e. MIC > 4 µg/mL) (Figure 6C) though, similar frequencies of trimethoprim resistance was observed for both species (Figure 6C). Thus, a difference in intrinsic evolvability under colistin selection was also reflected in the frequency of resistance among clinical strains. Since this interpretation could be confounded by relatively small sample size or local patterns of colistin use in clinical practice, we asked if similar trends were observed for isolates from around the world. To address this question, we collated MIC values of 2558 *E. coli* isolates and 1811 *K. pneumoniae* isolates of human/animal origin from 36 published studies across 20 countries (Supplementary File 4). Here too, *K. pneumoniae* isolates had higher colistin MICs (Figure 6D). We also looked at mechanisms of resistance in these isolates, wherever available (Supplementary File 4). While 94% of resistant *K. pneumoniae* harboured mutations in either BasSR or PhoQP pathways, only 18% of resistant *E. coli* showed mutations in these genes (Figure 6E). Instead, 82% of resistant *E. coli* isolates harboured MCR-genes, while only 6% of resistance isolates of *K. pneumoniae* were MCR-positive (Figure 6E). These results demonstrated that divergent genotype-phenotype maps and the resulting differences in intrinsic evolvability translated to different frequencies and genetic mechanisms of colistin resistance in clinical strains of *E. coli* and *K. pneumoniae* as well.

## Discussion

Genes that confer antimicrobial resistance in bacteria are identified by their strength of association with a resistant phenotype, as well as mechanistic or functional relation to the antibiotic target (exemplified by [80–82]). These genes, referred to here as “effectors of resistance”, are expected to have the largest contribution towards resistance in terms of MIC. As a result, it is reasonable to expect that conservation of effectors among related bacteria implies a similar capacity for resistance evolution. In this study, we demonstrate that this extrapolation is simplistic, and requires detailed experimental validation for individual bacterial pathogens. Our work shows that even when effector genes are conserved, genotype-phenotype maps of adaptive mutations can diverge substantially, such as for colistin resistance between two genera of Enterobacteriaceae, i.e. *Escherichia* and *Klebsiella*. This divergence was not due to the proximate impact of adaptive mutations on protein function, rather resulted from molecular-to- organism-level translation of phenotypic effects. Importantly, divergent genotype-phenotype maps directly affected evolvability under antibiotic pressure, as well as genetic strategies for resistance among clinical strains of *E. coli* and *K. pneumoniae*. This concordance between laboratory and clinical data shed light on how evolvability as a trait varies across related bacterial species. Evolvability, when viewed as ‘speed’ of adaptation is known to differ between different bacterial populations and environments [83, 84]. On the other hand, evolvability as ‘extent’ of adaptation, more relevant to the present study, is expected to be similar among closely related species facing stringent selection pressures like high antibiotic concentrations. Our study challenges this expectation and demonstrates that predictability in the evolution of drug resistance is confounded by divergences in genotype-phenotype maps.

A key difference between the two antibiotics used in our study is their molecular target. Trimethoprim is a substrate-mimicking enzyme inhibitor [85], while colistin binds to and disrupts the membranes of Gram-negative bacteria [86]. As a result, trimethoprim directly selected drug-target mutations, while colistin selected regulatory mutations that altered the expression of cell-envelope modifying enzymes. It is tempting to extrapolate our key conclusions regarding conservation of adaptive trajectories more generally to “protein targeting” and “non-protein targeting” antibiotics. Indeed, high concordance in target-site mutations even among distantly related organisms have been observed for other protein-targeting antibiotics as well, like rifampicin, which inhibits the beta-subunit of RNA polymerase [14, 87–89]. In part, this may be because of high sequence and structural conservation of essential proteins like DHFR or RpoB across bacteria. However, even when genotype-phenotype maps of drug-target mutations were similar, such as for trimethoprim in our study, we note that target mutations alone had a limited ability to explain resistance quantitatively at later time points of evolution. An increasing accumulation of non-target mutations, which independently led to only mild changes in drug MIC/IC50, was responsible for high resistance levels in conjunction with DHFR mutations. These results support the view that antibiotic resistance is a polygenic trait [90], and predicting resistance using single or small number of loci requires caution. As an example from the present study, a jackpot mutation combination in DHFR alone would not be classified as resistant according to the CLSI breakpoint for trimethoprim resistance (*E. coli* folA-D27E+F153S: MIC = 9.85±1.87 μg/mL, CLSI breakpoint ≥16 μg/mL), unless supported by non-target mutations. Such discrepancies have important repercussions for sequence-based diagnostics for AMR in bacterial pathogens, as highlighted by Grad et al. in a recent commentary [91]. Another confounding factor from our experiments was antibiotic efflux, which was enhanced under extended duration of drug pressure possibly as a result of genetic and phenotypic adaptation. The role of phenotypic adaptations such as efflux or biofilm induction in bacteria has been repeatedly discussed in different contexts such as reversible resistance, tolerance or persistence [92–95]. We propose that the interaction between phenotypic and genetic resistance strategies requires greater investigation to more accurately predict the phenotypes of resistant bacteria.

Though clinical breakpoints, such as CLSI or EUCAST, are useful for classifying bacteria as resistant or sensitive in clinical practice, our results for evolvability under colistin pressure highlight an important short-coming of this framework. Ancestrally, *E. coli* and *K. aerogenes* are both colistin-sensitive (CLSI breakpoint for resistance MIC = 4 μg/mL [52]), and colistin is expected to be a strong therapeutic candidate for both kinds of infections. However, since the capacity for resistance evolution based on chromosomal mutations was different, the long-term utility of colistin against these two species is expected to differ dramatically as well. Indeed, our meta-analyses of MIC values for clinical *E. coli* and *K. pneumoniae* isolates mirrored the intrinsic differences in evolvability measured under laboratory conditions. Another important outcome of dissimilar evolvabilities was the greater reliance of *E. coli* on plasmid-borne colistin resistance (*mcr* genes). Earlier work from the MacLean lab has shown that the pre-existence of MCR-1 greatly facilitated adaptation of *E. coli* under colistin pressure [96], supporting our claim. Other studies in *E. coli* also show that genomically encoded EptA is a poor substitute for MCR-1, possibly due to their different fitness costs and biochemical activities [51, 97]. We note, however, that under-representation of *mcr* genes in *Klebsiella* may also be due to other factors like fitness costs of horizontally transferred genes and constrains on plasmid compatibility [98, 99]. Regardless, we highlight a need to analyse the capacity for adaptation across individual pathogen-drug pairs to understand long term utility of antibiotic therapies for specific infections.

In summary, the results presented here throw up important considerations for how fundamental evolutionary genetics can contribute to a better understanding of the evolution of AMR. Firstly, defining intrinsic evolvabilities of pathogens using experimental pipelines for existing and new drugs may provide a set of guidelines for clinical use of broad spectrum antibiotics. At the very least, we expect that it will provide a much-needed framework to decide which infections broad-spectrum antibiotics should be deployed against. Second, the clear definition of species-specific genotype-phenotype maps for resistance are of vital importance. Beyond predicting the success of treatment, robust genotype-phenotype maps form the foundation of molecular diagnostics for AMR. We suggest caution in directly extrapolating genotype-phenotype relationships across bacteria, as even closely related species may respond differently to the same antibiotic over evolutionary time scales. Finally, resistance to antibiotics needs to be viewed as a quantitative phenotype that is shaped by interactions between genetic and non-genetic adaptations. We propose that the intrinsic evolvability of bacterial strains depends on both kinds of adaptive mechanisms, and the role of horizontally acquired genes that confer resistance should be viewed in light of these intrinsic capacities.

## Methods

### Bacterial strains and culture conditions

*E. coli* K-12 MG1655 and *K. aerogenes* ATCC 13048, and their derivatives, were cultured in Luria-Bertani broth (LB, Himedia, India) or grown as colonies on Luria-Bertani agar (LA, Himedia, India). Growth media was supplemented with trimethoprim (Sigma-Aldrich, Germany) or colistin (Himedia, India) as required. The following antibiotics were added to the media for selection of knockouts and transformants: chloramphenicol (Chl, 30 μg/mL), ampicillin (Amp, 100 μg/mL), kanamycin (Kan, 30 μg/mL), gentamicin (Gent, 25 μg/mL) and streptomycin (Strep, 50 μg/mL). Broth cultures were incubated at 37°C with shaking at 180 rpm and plates were incubated at 37°C. Glycerol stocks for freezing bacterial strains and populations were prepared by adding sterile 50% glycerol to saturated overnight cultures in a 1:1 ratio and stored at −80°C.

The strains used in this study are as follows:

| Sr. No. | Strain name | Description | Source/Reference |
| --- | --- | --- | --- |
| 1. | *E. coli* K-12 MG1655 | Wild-type *E. coli* used as ancestor for evolution experiments and as a reference strain for phenotypic and genotypic comparisons. | Available in the laboratory |
| 2. | *K. aerogenes* ATCC 13048 | Wild-type *K. aerogenes* used as ancestor for evolution experiments and as a reference strain for phenotypic and genotypic comparisons. | American Type Culture Collection (ATCC) |
| 3. | *E. coli* $\Delta lon$ | Isogenic knockout of *lon* gene ($\Delta lon::kan$) | Matange (2020) [30] |
| 4. | *E. coli-folA-F153S* | Scarless genomic knock-in harbouring Phe153Ser mutation in *folA* | This study |
| 5. | *E. coli-folA-D27E-F153S* | Scarless genomic knock-in harbouring Asp27Glu and Phe153Ser mutations in *folA* | This study |
| 6. | *E. coli-folA-D27E-F153S $\Delta acrR$* | Scarless genomic knock-in harbouring Asp27Glu and Phe153Ser mutations in *folA* and a knockout of *acrR* gene ($\Delta acrR::kan$) | This study |
| 7. | *E. coli-folA*-<br>D27E-F153S<br>$\Delta mgrB$ | Scarless genomic knock-in harbouring<br>Asp27Glu and Phe153Ser mutations in *folA* and<br>a knockout of *mgrB* gene ($\Delta mgrB::kan$) | This study |
| 8. | *E. coli-folA*-<br>D27E-F153S<br>$\Delta rng$ | Scarless genomic knock-in harbouring<br>Asp27Glu and Phe153Ser mutations in *folA* and<br>a knockout of *rng* gene ($\Delta rng::kan$) | This study |
| 9. | *E. coli-folA*-<br>D27E-F153S<br>$\Delta acrR \Delta rng$ | Scarless genomic knock-in harbouring<br>Asp27Glu and Phe153Ser mutations in *folA* and<br>knockouts of *acrR* and *rng* genes ($\Delta acrR::kan$<br>$\Delta rng::FRT$) | This study |
| 10. | *E. coli-folA*-<br>D27E-F153S<br>$\Delta mgrB \Delta acrR$ | Scarless genomic knock-in harbouring<br>Asp27Glu and Phe153Ser mutations in *folA* and<br>knockouts of *mgrB* and *acrR* genes ($\Delta acrR::kan$<br>$\Delta mgrB::FRT$) | This study |
| 11. | *E. coli-folA*-<br>D27E-F153S<br>$\Delta rng \Delta mgrB$ | Scarless genomic knock-in harbouring<br>Asp27Glu and Phe153Ser mutations in *folA* and<br>knockouts of *rng* and *mgrB* genes ($\Delta rng::kan$<br>$\Delta mgrB::FRT$) | This study |
| 12. | *E. coli-folA*-<br>D27E-F153S<br>$\Delta rng \Delta acrR$<br>$\Delta mgrB$ | Scarless genomic knock-in harbouring<br>Asp27Glu and Phe153Ser mutations in *folA* and<br>knockouts of *acrR*, *rng* and *mgrB* genes<br>($\Delta acrR::kan \Delta rng::FRT \Delta mgrB::FRT$) | This study |
| 13. | *E. coli \Delta acrR* | Isogenic knockout of *acrR* gene ($\Delta acrR::kan$) | This study |
| 14. | *E. coli \Delta mgrB* | Isogenic knockout of *mgrB* gene ($\Delta mgrB::kan$) | [50] |
| 15. | *E. coli \Delta rng* | Isogenic knockout of *rng* gene ($\Delta rng::kan$) | This study |
| 16. | *E. coli \Delta acrR*<br>$\Delta rng$ | Double knockout of *acrR* and *rng*<br>($\Delta acrR::kan \Delta rng::FRT$) | This study |
| 17. | *E. coli \Delta mgrB*<br>$\Delta acrR$ | Double knockout of *mgrB* and *acrR*<br>($\Delta acrR::kan \Delta mgrB::FRT$) | This study |
| 18. | *E. coli* $\Delta rng$<br>$\Delta mgrB$ | Double knockout of *rng* and *mgrB*<br>($\Delta rng::kan \Delta mgrB::FRT$) | This study |
| 19. | *E. coli* $\Delta acrR$<br>$\Delta rng \Delta mgrB$ | Triple knockout of *acrR*, *rng* and *mgrB*<br>($\Delta acrR::kan \Delta rng::FRT \Delta mgrB::FRT$) | This study |
| 20. | *E. coli* DH10B | *E. coli* strain used for cloning and genetic manipulations | Available in the laboratory |
| 21. | *E. coli* CC118 | *E. coli* strain for propagation and manipulation of pSEVA612S plasmid, which contains an R6K origin of replication | Kind gift from Prof. Gad Frankel, Imperial College, London, U.K. |
| 22. | *E. coli* pRK2013 helper strain | *E. coli* helper Strain for triparental conjugation | Kind gift from Prof. Gad Frankel, Imperial College, London, U.K. |
| 23. | *E. coli* $\Delta basS$ | Isogenic knockout of *basS* gene ($\Delta basS::kan$) | This study |
| 24. | *E. coli* $\Delta phoQ$ | Isogenic knockout of *phoQ* gene ($\Delta phoQ::kan$) | [50] |
| 25. | *E. coli* $\Delta arnT$ | Isogenic knockout of *arnT* gene ($\Delta arnT::kan$) | This study |
| 26. | *E. coli* $\Delta arnB-F$ | Isogenic knockout of the *arn* operon (*arnBCADTEF*) ($\Delta arnB-F::kan$) | This study |
| 27. | *E. coli* $\Delta eptA$ | Isogenic knockout of *eptA* gene ($\Delta eptA::kan$) | This study |
| 28. | *E. coli* Cft073 | Uropathogenic *E. coli* strain used as ancestor for evolution experiments | Kind gift from Prof. Gad Frankel, Imperial College, London, U.K. |
| 29. | *E. coli* S34 | *E. coli* isolated from river water used as ancestor for evolution experiments | This study |

List of plasmids used in this study:

| Sr. No. | Plasmid | Description | Source/Reference |
| --- | --- | --- | --- |
| 1. | pPRO-*folA* | *E. coli folA* gene cloned into pPRO-Ex-Htb plasmid for heterologous expression, Amp<sup>R</sup> | Matange et al. (2018) [46] |
| 2. | pPRO-*folA*-I94L | Derivative of pPRO-*folA* expressing Ile94Leu mutant of DHFR, Amp<sup>R</sup> | Matange (2020) [30] |
| 3. | pPRO-*folA*-D27E | Derivative of pPRO-*folA* expressing Asp27Glu mutant of DHFR, Amp<sup>R</sup> | This study |
| 4. | pPRO-*folA*-F153S | Derivative of pPRO-*folA* expressing Phe153Ser mutant of DHFR, Amp<sup>R</sup> | This study |
| 5. | pPRO-*folA*-A26T | Derivative of pPRO-*folA* expressing Ala26Thr mutant of DHFR, Amp<sup>R</sup> | This study |
| 6. | pPRO-*folA*-D27E-F153S | Derivative of pPRO-*folA* expressing Asp27Glu and Phe153Ser double mutant of DHFR, Amp<sup>R</sup> | This study |
| 7. | pPRO-*folA*-I94L-A26T | Derivative of pPRO-*folA* expressing Ile94Leu and Ala26Thr double mutant of DHFR, Amp<sup>R</sup> | This study |
| 8. | pPRO-*folA*-F153S-I94L | Derivative of pPRO-*folA* expressing Phe153Ser and Ile94Leu double mutant of DHFR, Amp<sup>R</sup> | This study |
| 9. | pPRO-*folA*-D27E-I94L | Derivative of pPRO-*folA* expressing Asp27Glu and Ile94Leu double mutant of DHFR, Amp<sup>R</sup> | This study |
| 10 | pPRO-*folA*-F153S-A26T | Derivative of pPRO-*folA* expressing Phe153Ser and Ala26Thr double mutant of DHFR, Amp<sup>R</sup> | This study |
| 11 | pPRO-*folA*-D27E-A26T | Derivative of pPRO-*folA* expressing Asp27Glu and Ala26Thr double mutant of DHFR, Amp<sup>R</sup> | This study |
| 12 | pSEVA612S-GFP | pSEVA612S plasmid encoding GFP and harbouring an R6K origin of replication as well as *oriT*, Gent<sup>R</sup> | Kind gift from Prof. Gad Frankel, Imperial College, London, U.K. |
| 13 | pSEVA612S-*folA* | pSEVA612S plasmid in which GFP is replaced by *folA*, Gent<sup>R</sup> | This study |
| 14 | pSEVA612S-*folA*-D27E | pSEVA612S plasmid encoding DHFR Asp27Glu, Gent<sup>R</sup> | This study |
| 15 | pSEVA612S-*folA*-F153S | pSEVA612S plasmid encoding DHFR Phe153Ser, Gent<sup>R</sup> | This study |
| 16 | pSEVA612S-*folA*-D27E-F153S | pSEVA612S plasmid encoding DHFR Asp27Glu and Phe153Ser, Gent<sup>R</sup> | This study |
| 17 | pACBSR | Temperature sensitive plasmid expressing I-*SceI* enzyme and lambda recombinase system under an arabinose inducible promoter, Strep<sup>R</sup> | Kind gift from Prof. Gad Frankel, Imperial College, London, U.K. |
| 18 | pCA24N-*basS* | pCA24N containing *basS* with an N-terminal hexa-histidine tag under the T5-lac promoter, Chl<sup>R</sup> | ASKA library [100] |
| 19 | pCA24N-*basS*-T156P | Plasmid pCA24N-*basS*, containing Thr156Pro mutation in *basS*, Chl<sup>R</sup> | This study |
| 20 | pCA24N-*phoQ* | pCA24N containing *phoQ* with an N-terminal hexa-histidine tag under the T5-lac promoter, Chl<sup>R</sup> | ASKA library [100] |
| 21 | pCA24N-*phoQ*-K353E | Plasmid pCA24N-*phoQ*, containing Lys353Glu mutation in *phoQ*, Chl<sup>R</sup> | This study |
| 22 | pCA24N-<br>*phoQ*-<br>L395Q | Plasmid pCA24N-*phoQ*, containing<br>Leu395Gln mutation in *phoQ*, Chl<sup>R</sup> | This study |
| 23 | pCA24N-<br>*phoQ*-<br>S288R | Plasmid pCA24N-*phoQ*, containing<br>Ser288Arg mutation in *phoQ*, Chl<sup>R</sup> | This study |
| 24 | pCA24N-<br>*phoQ*-V42M | Plasmid pCA24N-*phoQ*, containing<br>Val42Met mutation in *phoQ*, Chl<sup>R</sup> | This study |
| 25 | pCA24N-<br>*phoQ*-<br>D436N | Plasmid pCA24N-*phoQ*, containing<br>Asp436Asn mutation in *phoQ*, Chl<sup>R</sup> | This study |
| 26 | pCA24N-<br>*arnT* | pCA24N containing *arnT* with an N-terminal hexa-histidine tag under the T5-lac<br>promoter, ChlR | ASKA library [100] |
| 27 | pCA24N-<br>*eptA* | pCA24N containing *eptA* with an N-terminal hexa-histidine tag under the T5-lac<br>promoter, ChlR | ASKA library [100] |
| 28 | pCA24N-KA*eptA* | pCA24N containing *eptA* from *K.*<br>*aerogenes* with an N-terminal hexa-histidine tag under the T5-lac promoter,<br>ChlR | This study |
| 29 | pGDP2-MCR-1 | *mcr-1* gene cloned into the low copy<br>number vector pGDP2 | Addgene plasmid # 118404<br>[101] |

### Sequence alignments, structural prediction and analysis

Pairwise sequence alignment of *E. coli* and *K. aerogenes* DHFR proteins was created with EMBOSS Needle using default parameters [102] and the alignment was visualized in Jalview [103].

Multiple sequence alignment of protein effectors of resistance was created with the help of Clustal Omega [102]. Jalview was used for visualisation of the alignment and calculation of percentage identities between sequences [103].

Structures of *E. coli* and *K. aerogenes* DHFR proteins were generated on the AlphaFold3 server [104] by using default parameters. Visualization of protein structures, structural alignment and analysis of average Cα-RMSD values was done with the help of UCSF ChimeraX [105].

### Characterising antibiotic resistance phenotypes

#### Microbroth dilution assay for determination of MIC and IC50

MIC and IC50 values were determined by using microbroth dilution assay in 96-well microtitre plates as described earlier [13, 106]. Briefly, primary cultures were set up in 1 mL drug-free LB and incubated overnight until saturation at 37°C. These primary cultures were then inoculated (0.1% for trimethoprim, 1% for all other antibiotics, unless specified) in a 96-well plate into 150 μL LB/well, in which a 2-fold dilution series of the antibiotic was set up. No antibiotic growth controls were kept for each strain. Plates were incubated at 37°C for 18 hours and OD_600_ was measured. Growth in the presence of antibiotic at each concentration was normalised to growth in the absence of antibiotic (set to 1). The data were then fitted to a nonlinear regression (log inhibitor versus response) curve using Graphpad Prism (version 9.1.4). Following equation was used to determine IC50 value. where Min: Minimum OD value; Max: Maximum OD value (=1); and Hill Slope: Slope of curve after fitting.

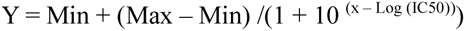

From this model, values of IC50, i.e., drug concentration needed to achieve 50% growth inhibition, were estimated. Fold changes were calculated where needed as a ratio of IC50 of test and reference strains. IC50s were used for analysing pairwise epistasis of DHFR mutations using the multiplicative model accounting for error propagation as described in Borrell et al. (2013) [107].

#### Epsilometry (E-tests)

Primary cultures were set up in 1 mL drug-free LB and incubated overnight until saturation at 37°C. An aliquot (200 μL) was spread on an LA plate. The plates were dried for 10 minutes and trimethoprim/colistin E-test strips (Himedia, India) were placed on the agar surface. Plates were incubated for 18 hours at 37°C and imaged.

#### Colistin-induced cell death

This assay was performed to test the viability of wild-type *E. coli* and *K. aerogenes* strains upon 6 hours of growth in varying concentrations of colistin. Primary cultures were set up in 1 mL drug-free LB and incubated overnight until saturation at 37°C. Secondary cultures were set up by transferring 30 μL of the primary culture in 3 mL LB containing 0, 1, 2, 4 and 8 μg/mL colistin. The cultures were incubated for 6 hours at 37°C with shaking (180 rpm). Cultures were then serially diluted (10-10^6^ fold) and 10 μL of each dilution was spotted onto LA plates. Upon drying, plates were incubated for 18 hours at 37°C, and viable counts were calculated as CFU/mL.

#### Colony formation on trimethoprim-supplemented solid media

A colony formation assay was performed to determine the susceptibility of *rng-*D268A to trimethoprim. As described in Balachandran et al. (2025) [26], primary cultures were set up in 1 mL-drug-free LB and incubated overnight until saturation at 37°C with shaking at 180 rpm. Strains transformed with pCA24N-*rng* and pCA24N-*rng*-D268A were also grown in the presence of 0.1 mM IPTG. Cultures were serially diluted using a 10-fold dilution scheme. An aliquot (10 μL) of each dilution was spotted onto LA plates containing 0.1mM of IPTG and varying trimethoprim concentrations. Plates were incubated overnight at 37°C, imaged and CFU/mL were estimated.

### Adaptive laboratory evolution of trimethoprim and colistin resistance

To compare evolution of resistance in *E. coli* and *K. aerogenes*, adaptive laboratory evolution was performed in presence of increasing concentrations of trimethoprim and colistin. Primary cultures were set up in 1 mL drug-free LB for *E. coli* K-12 MG1655 and *K. aerogenes* ATCC 13048 and incubated at 37°C overnight. From this primary culture, 5 replicate lineages of each strain were set up by inoculating 10 μL of the culture in 1 mL LB supplemented with 100 ng/mL trimethoprim or colistin (∼0.1X MIC). The cultures were incubated at 37°C overnight with shaking at 180 rpm, and this tube was labelled as Passage 1 (∼7 generations). From this culture, 10 μL was transferred again into fresh LB supplemented with antibiotic until 30 passages (∼210 generations), with a 5X increase in antibiotic concentration every 5 passages (∼35 generations). An aliquot of the evolving populations (500 μL) was frozen every 5 passages, i.e. before increasing antibiotic concentration. Lack of turbidity in media after 24 hours of incubation at any passage during evolution was inferred as extinction.

### Whole genome sequencing of evolved populations and mutation calling

Population whole genome sequencing and mutation calling were performed using the same methodology as described in Vinchhi and Yelpure et al. (2023) [13] and Balachandran et al. (2025) [26]. Briefly, glycerol stocks of evolving populations were revived in 3 mL LB and incubated overnight. Genomic DNA was extracted from bacterial pellets using spin columns (Himedia, India). Library preparation and sequencing services were provided by Eurofins India Pvt Ltd. Paired-end next generation sequencing was performed for the genomic DNA on a MiSeq system (Illumina, USA) with an average read length of 150 bps. Alignment and mutation calling was performed using Breseq [108]. Genomes of *E. coli* K-12 MG1655 (NCBI Accession: U00096), *Klebsiella aerogenes* ATCC 13048 (acquired from ATCC), *Escherichia coli* CFT073 (NCBI Accession: NZ_CP051263) and genome assembly of *E. coli* Isolate S34 (PRJNA1491519) were used as references for aligning reads using bowtie2. Sequencing data were analysed in the population mode using default parameters [108] and mutations and new junctions in the genomes of evolved populations were identified. All mutations and new junctions present in the respective ancestor were excluded. Mutations which occurred with a frequency of 10% or higher in the population were considered for further analysis. Raw sequencing data is available on Genbank (PRJNA1489306, PRJNA1489700, PRJNA1489791, PRJNA1493335, PRJNA1494217).

### Site-directed mutagenesis for generation of point mutations in heterologously expressed DHFR, BasS, PhoQ and Rng

PCR-based site-directed mutagenesis was used to generate point mutations in pCA24N-*phoQ*, pCA24N-*basS*, pCA24N-*rng* and pPRO-*folA* using the protocol described by Shenoy and Visweswariah (2003)[109]. A slight modification was made to this protocol by setting up 20 μL PCR reactions with both forward and reverse mutagenic primers, except for generation of pPRO-folA-A26T, where a single primer was used. The PCR product was digested using *Dpn*I and transformed into *E. coli* DH10B. Plasmids were isolated and mutants were confirmed by restriction digestion and Sanger sequencing (Barcode BioSciences, India). Double mutants were generated by using single mutants as the template. However, pPRO-*folA*-A26T-D27E was generated by introducing both mutations using a single pair of primers. Mutagenic primers used in this study are listed in Supplementary file 5.

### Generation of scarless genomic *folA* knock-in strains of *E. coli*

Scarless genomic knock-ins (*E. coli*-*folA*-F153S and *E. coli*-*folA*-D27E-F153S) were generated using triparental conjugation as described in Low et al. (2022) [110] in three steps.

#### Step 1: Generation of pSEVA-folA clones

In the first step, *folA* was amplified from *E. coli* K-12 MG1655 genomic DNA, along with 5’- and 3’-homology regions (500 bp each) using the folA_homology_fwd and folA_homology_rev primers (Supplementary file 5).

The PCR product was cloned into pSEVA612S plasmid by Gibson assembly to yield pSEVA612S-*folA*. The generated pSEVA612S-*folA* plasmid was used as a template to generate pSEVA612S-*folA*-D27E and pSEVA612S-*folA* F153S, by amplifying the *folA* gene using folA_homology primers and appropriate *folA* mutagenic primers in two parts, and then performing a three-part Gibson assembly (Part 1: amplified from pSEVA612S-folA by using folA_homology_fwd and folA_mutant(D27E/F153S)_rev; Part 2: amplified from pSEVA612S-folA as a template by using folA_homology_rev and folA_mutant(D27E/F153S)_fwd Part 3: pSEVA612S linearized with the I-*Sce*I primer using pSEVA612S-GFP as the template). The primers used for these PCRs are listed in Supplementary file 5.

Double mutant pSEVA612S -*folA*-D27E-F153S was generated by using pSEVA612S-folA-D27E as the template and folA_F153S mutagenic primers. For the three-part assembly, purified fragments were combined in a 5:1 insert:vector molar ratio, and incubated with Gibson Assembly Master Mix (New England Biolabs, USA) at 50°C for 30 minutes. The reaction was then transformed into chemically competent *E. coli* CC118 and transformants were selected using gentamicin (25 μg/mL). All generated clones were confirmed by restriction digestion and Sanger sequencing (Genematrix, India).

#### Step II: Triparental conjugation and generation of single cross-over merodiploid strains

For triparental conjugation, an aliquot (20 μL) of an overnight culture of *E. coli* CC118 containing the appropriate mutant pSEVA612S-folA plasmid was spotted on an LA plate and allowed to completely dry. Next, 20 μL of overnight culture of *E. coli* pRK2013 helper strain was spotted over the donor strain and allowed to dry. The plate was then incubated at 37°C for 2 hours. Next, 40 μL of overnight culture of the recipient strain, *E. coli* K-12 MG1655 transformed with pACBSR was spotted, dried and incubated for 4 hours at 37°C. The growth obtained was scraped using a sterile nichrome loop and resuspended in 1 mL LB. An aliquot (100 μL) of the bacterial suspension was then plated on an LA plate supplemented with gentamicin (25 μg/mL) and streptomycin (50 μg/mL). The plate was incubated for 18 hours at 37°C to select for single crossover merodiploid colonies.

#### Step III: Generation of scarless knock-in strains from merodiploid single cross-overs

Three merodiploid colonies were selected at random and inoculated in 3 mL LB containing streptomycin (50 μg/mL) and L-arabinose (0.4%) to induce the expression of I-*Sce*I enzyme and lambda red recombination system present on pACBSR. After induction at 37°C for 2 hours, the cultures were streaked on an LA plate to obtain double cross-over strain colonies. Colonies (10-20) were first screened phenotypically for gentamicin sensitivity (25 μg/mL) and trimethoprim resistance (1 μg/mL). Putative scarless knock-ins were also screened using colony PCR with *folA*-homology primers followed by restriction digestion, and finally confirmed by Sanger sequencing of the *folA* locus (Genematrix, India). Confirmed knock-in strains were passaged in 3 mL drug-free LB for five days, with 1% of the culture being transferred to fresh LB on each day to cure pACBSR.

### Generation of single and multiple gene knockouts in *E. coli*

All gene knockouts, except for a knockout of the full *arn* operon, were generated in *E. coli* using P1 transduction, by moving the kanamycin resistance cassette-marked gene deletions from the respective strains in the Keio collection[111] into *E. coli* K-12 MG1655. Colony PCRs using gene-specific primers were used to confirm transductants. Double and triple knockouts were generated in *E. coli* K-12 MG1655 by first removing the kanamycin cassette from the single/double knockout using pCP20 plasmid expressing FLP recombinase [112] and by introducing the second/third knockout allele via P1 transduction.

Knockout of the entire *arn* operon (*arnBCADTEF*) in *E. coli* was generated by using lambda red recombination system [111]. Primers pKD3,4_arnB_fwd and pKD3,4_arnF_rev, containing the 5’- and 3’-homology regions of *arnB* and *arnF* genes, respectively, were used to amplify the kanamycin and chloramphenicol cassettes from pKD4 and pKD3 by PCR (Supplementary file 5). The PCR product was purified and electroporated into competent cells of *E. coli* K-12 MG1655 containing the red helper plasmid pKD46. Knockouts were confirmed by using gene-specific primers.

### Cloning of *K. aerogenes eptA* into pCA24N for heterologous expression in *E. coli*

To check for the effect of overexpression of *eptA* from *K. aerogenes* in *E. coli*, *eptA* gene from *K. aerogenes* was cloned into pCA24N using restriction enzyme-based cut-paste cloning. The *eptA* gene was amplified from *K. aerogenes* genomic DNA using primers pCA24N_KA_eptA_SfiI_fwd and pCA24N_KA_eptA_HindIII_rev (Supplementary file 5), and inserted into the *Sfi*I and *Hin*dIII sites of pCA24N in frame with an N-terminal hexa-his tag. Clones obtained after ligation were confirmed by Sanger sequencing (Genematrix, India).

### Monitoring efflux activity using Hoechst H33342 accumulation assay

Accumulation of Hoechst H33342 dye was used to measure efflux activity in *E. coli* and *K. aerogenes* as described in Coldham et al. (2010) [113] with some modifications. Saturated cultures were first sub-cultured by inoculating 30 μL in 3 mL drug-free LB, and incubated for 3 hours at 37°C. Post incubation, cells were pelleted by centrifugation at 13,000 rpm for 1 minute and resuspended in 1 mL PBS. OD_600_ of all cultures was then adjusted to 0.1. Aliquots of the bacterial suspension in PBS (180 μL) was added to a 96 well microtitre plate, and 20 μL of 100 μM bisbenzimide H33342 trihydrochloride (Sigma-Aldrich, Germany) was added to the culture, resulting in a final concentration of 10 μM of the dye. Fluorescence was measured (excitation filter: 355 nm, emission filter: 460 nm, 5 flashes per well) using a multimode plate reader (Revvity, USA) every 1 minute for 150 minutes at 25°C.

### Transcriptome analyses using RNA sequencing

RNA sequencing was performed to analyse the transcriptome of *E. coli* Δ*rng*. Triplicate primary cultures of *E. coli* wild type and Δ*rng* strains were grown in 3 mL drug-free LB overnight until saturation. On the next day, 30 μL was inoculated in 3 mL drug-free LB in triplicates and incubated at 37°C with shaking at 180 rpm for 3 hours. All cultures were pelleted by centrifugation at 13,000 rpm for 1 minute. Pellets were resuspended in 1 mL of RNAlater (Invitrogen, U.S.A). Total RNA was extracted using TRIzol Reagent (Invitrogen, USA), and the quality of the extracted RNA was estimated using TapeStation (Agilent, USA). Ribosomal RNA was depleted using the Ribozero kit. This was followed by library preparation and RNA-sequencing performed on Illumina Miseq platform with an average read length of 150 bp. Processed reads were aligned to *E. coli*’s reference genome (NCBI Accession: NC_00096.3) using STAR (v2.7.10a). Abundance estimation was performed for counting reads corresponding to genes, promoter regions, and genomic bins using featureCounts (1.34.0). RNA extraction, ribodepletion, library preparation, sequencing and preliminary analyses were performed by Eurofins Genomics, India. Differential gene expression analysis was performed using DESeq2. All strains were compared to the wild type to obtain the Log_2_(fold-change) values and statistical significance. Gene names were inferred from the Ecocyc database[114]. Raw sequencing reads are available in Genbank (PRJNA1399355).

### Gene-specific quantitative RT-PCR

RNA isolation and quantitative PCRs were performed as described in Vinchhi and Yelpure et al. (2023) [13]. Briefly, for RNA isolation, bacterial cultures were grown for 3 hours in drug-free LB and pelleted by centrifugation. Total RNA was extracted using TRIzol Reagent (Invitrogen, USA) and quantification for was done spectrophotometrically. Evaluation of RNA quality was done by electrophoresis on a 1% agarose gel. Extracted RNA (20 μg) was treated with RNAse-free recombinant DNase I (Takara, Japan), followed by cleanup using the NucleoSpin RNA clean-up columns kit (MACHEREY-NAGEL, Germany). Cleaned-up RNA (1.5 μg) was used for the reverse transcription reaction, using PrimeScript RT Reagent Kit (Takara, Japan). For the RT reaction, buffering temperature was 25°C, for 2 minutes, and cDNA synthesis was carried out at 37°C for 30 minutes. This was followed by heat inactivation of the RT enzyme at 85°C for 90 seconds. Prepared cDNA and -RT controls were serially diluted by 10, 100 and 1000 fold in nuclease-free waterand stored at −20°C until further use. Semi-quantitative PCRs were performed before qPCRs using gene-specific primers (Supplementary file 5) to ensure complete degradation of residual genomic DNA, and to optimise PCR conditions such as primer concentration.

Quantitative RT-PCR (qPCR) was performed on Bio-Rad CFX96 Real-Time System (Bio-Rad, USA) using a two-step protocol with an initial denaturation at 95°C for 30 seconds, followed by 40 cycles of denaturation at 95°C for 5 seconds and annealing/extension at 60°C for 30 seconds. Melt curve analysis was performed after 40 cycles. Each qPCR reaction was 20 μL and contained the following: 10 μL of 2X TB GreenPremix Ex Taq II (Tli RNaseH Plus) (Takara, Japan), 0.5 μL each of gene-specific forward and reverse primers (10 μM stock, Supplementary file 5), 8 µL of nuclease-free water, and 1 µL of the appropriate dilution of cDNA. Known concentrations of genomic DNA from *E. coli* and *K. aerogenes* were used to construct a standard graph for each gene-specific primer and quantitate expression level. 16S rRNA was used as a normalizing control for cDNA from *E. coli* and *rpoD* was used as a normalizing control for cDNA from *K. aerogenes*. These controls were used to calculate fold changes in gene expression with respect to the wild type using the standard graph method.

### NPN uptake assay for outer membrane permeability after colistin treatment

NPN uptake assay was performed as described by Helander and Mattila-Sandholm (2000) [115]. Briefly, the bacterial strains were inoculated in 1 mL drug-free LB and incubated overnight until saturation. Cells were pelleted by centrifugation at 13,000 rpm at room temperature for 1 minute. The pellet was washed thrice with PBS and diluted to OD_600_ of 0.5 in PBS. An aliquot (100 μL) of this suspension was added in a 96-well microtitre plate containing 50 μL PBS. 1-N-phenylnaphthylamine (NPN, Sigma Aldrich, Germany) solution (50 μL of 40 μmol^−1^ prepared in PBS) was added to all samples. Additionally, the following controls were set up: 1. 200 μL PBS 2. 150 μL PBS with 50 μL NPN 3. 100 μL culture with 100 μL PBS. For measuring outer membrane permeability upon colistin exposure (1 μg/mL), 50 μL of 4 μg/mL colistin in PBS was used instead of buffer alone. The 96-well microtitre plate was kept in dark at room temperature for 1 hour and fluorescence was measured post incubation (excitation wavelength: 355 nm, emission wavelength: 405 nm) using a multimode plate reader (Revvity, USA). Fluorescence values were used to calculate the NPN uptake factor in presence as well as absence of colistin as given below.

NPN uptake factor = (Fluorescence of sample with NPN-Fluorescence of sample without NPN) ÷ (Fluorescence of PBS with NPN-Fluorescence of PBS without NPN)

### Propidium iodide assay for cell membrane disruption by colistin

This assay was adapted from a study by Sabnis et al. (2021) [45] to estimate inner membrane damage induced by colistin. Bacterial strains were inoculated in 1 mL drug free LB and incubated overnight at 37°C until saturation. A secondary culture was set up for each strain by inoculating 200 μL of the primary culture in 3 mL drug-free LB. To test the effect of colistin on membrane permeability, secondary cultures of all strains were also set up with LB supplemented with 1 μg/mL colistin. Cultures were incubated at 37°C for one hour and then centrifuged at 13000 rpm for 1 minute. The bacterial pellet was resuspended in PBS (to an OD_600_= 0.5). An aliquot (200 μL) of this suspension was transferred in a 96-well microtitre plate and 50 μL Propidium iodide (Sigma-Aldrich, Germany) was added to each well at a final concentration of 2.5 μM. PBS alone and no propidium iodide controls were kept for each strain. The plate was incubated for 3 hours at 25°C, and fluorescence (excitation wavelength: 535 nm, emission wavelength: 617 nm) and OD_600_ were measured using a multimode plate reader (Revvity, USA). Fluorescence values were corrected for cell number using OD_600_.

### Meta-analysis of resistance and mutations in colistin resistant clinical isolates of *E. coli* and *K. pneumoniae*

Trimethoprim and colistin susceptibilities of *E. coli* and *K. pneumoniae* clinical isolates collected at hospitals from North and West India were obtained from the work of Gheewalla et al. (2026) [79].

For the meta-analysis of colistin resistant *E. coli* and *K. pneumoniae* clinical isolates from around the world, a literature search was conducted using the following keywords in various combinations: ‘colistin resistance’, ‘*Escherichia coli’*, ‘*Klebsiella pneumoniae’*, ‘colistin MIC’ and ‘mutations’. Any study which either characterised colistin MICs or mechanisms of colistin resistance in either *E. coli* or *K. pneumoniae* of clinical origin were taken forward for the analysis. The following data were obtained from the selected studies: 1. year of publication, 2. region of study, 3. sample origin (human/animal origin), 4. number of isolates of *E. coli* or *K. pneumoniae* studied, 5. colistin MICs of all isolates, 6. number of isolates with *mcr* genes, 7. number of isolates with chromosomal mutations (in *phoQP*/*basSR*/*mgrB*). The papers selected for the meta-analysis as well as the compiled data are summarised in Supplementary File 4.

## Acknowledgement

We thank Prof. Gad Frankel from Imperial College (London) for sharing the plasmid system required for generating scarless genomic knock-in strains using triparental conjugation.

